# Vitamin D regulation of a SOD1-to-SOD2 antioxidative switch to prevent osteosarcoma transformation

**DOI:** 10.1101/2020.03.03.975854

**Authors:** Thomas S. Lisse

## Abstract

Superoxide, a form of reactive oxygen species (ROS), is catabolized by superoxide dismutase (SOD) and contributes to carcinogenesis via the oxidative damage it inflicts on cells. The aim of this research was to analyze the potential vitamin D-mediated regulation of the antioxidative “SOD1-to-SOD2 switch” within the human MG-63 osteosarcoma model. For this study, real-time PCR analysis was performed using MG-63 cells exposed to metabolically active 1,25(OH)_2_D_3_. Frist, a sustained statistically significant >2-fold suppression of proliferating cell nuclear antigen (PCNA) transcripts was observed after 10nM but not at 100nM of 1,25(OH)_2_D_3_ treatment, suggesting a cytostatic effect. In order to assess regulators of mitochondrial oxidative phosphorylation, gene expression of *COX2* and *COX4l1* of the mitochondrial complex IV and antioxidative enzymes (*SOD1, SOD2* and *Catalase* (*CAT*)) were monitored. For COX2 and COX4l1, no changes in gene expression were observed. However, a concomitant decrease in CAT and SOD1 mRNA, and increase in SOD2 mRNA after 24 hours of 10nM 1,25(OH)_2_D_3_ treatment were observed. A ~8-fold increase in SOD2 mRNA was apparent after 48 hours. The significant increase in SOD2 activity in the presence of vitamin D indicates an antioxidant potential and sensitization of vitamin D during osteosarcoma transformation and mitochondrial detoxification over time.

## Introduction

Vitamin D is a steroid hormone produced in the skin, or obtained in varying quantities through dietary or supplementary means^1^. Vitamin D is metabolized through two subsequent hydroxylation steps, resulting in its metabolically active form, 1,25(OH)_2_D_3_ (also known as calcitriol or 1,25D_3_). 1,25D_3_ exerts its genomic effects by binding to the vitamin D receptor (VDR), a member of the nuclear receptor super family ^2–13^. It has long been recognized that vitamin D has noncalcemic functions in controlling the cell cycle, differentiation and apoptosis, and its analogues are used in cancer prevention and treatment^14,15^. Understanding how vitamin D can regulate these processes is central to applications toward specialized cancer treatment^15^.

It is widely accepted that oxidative damage is one among several key factors that limits lifespan. Superoxide (O_2_^-^) anions are produced as a free radical reactive oxygen species (ROS) byproduct of oxygen metabolism and mitochondrial respiration, and can damage cells at high concentrations if left unchecked^16^. Superoxide anions are produced and accumulate in the matrix and the intermembrane space of the mitochondria, and are also produced from the enzymes NOX/DUOX localized at the plasma membrane^17^. Excessive superoxide in the cell results in damage to mitochondrial DNA, proteins and lipids, and ultimately leads to irreversible damage to organelles. To counter the effects of elevated superoxide anions, cells express superoxide-scavenging enzymes superoxide dismutase (SOD)^18^. SOD catalyzes the disproportionation of superoxide into either molecular oxygen or hydrogen peroxide as part of its antioxidant defense mechanism. Hence, SOD enzymes maintain a range of ROS in order to control potential toxicity, as well as downstream cellular and signaling functions^19^. There are three forms of SODs, and the localization of the SODs reflect their antioxidative roles within cells. SOD1 is mostly sequestered in the cytosol but can also localize to the mitochondria, SOD2 in the matrix of mitochondria^20^, and SOD3 is in the extracellular space^18^. Several regulators of SODs have been characterized. For example, SOD2 is regulated by SIRT3-mediated deacetylation that can attenuate excessive production of mitochondrial ROS under disease conditions^21^.

Vitamin D is known to exert its anti-cancer effects by a variety of mechanisms, most prominently studied is through G_1_ cell cycle arrest^7,10,22–24^. However, there are several studies that have looked at the role of vitamin D on ROS production and detoxification. For example, a recent paper has shown that supraphysiological levels of calcitriol exerts its anti-tumor effects in a mouse cell model of osteosarcoma via induction of the endoplasmic reticulum stress response with a concomitant increase in intracellular ROS^25^. In addition, cDNA microarray analysis using human prostate epithelial cells revealed that calcitriol (50nM) can promote SOD2 expression (2.01 fold increase, 24 hours)^26^, as well as the early expression of the antioxidant thioredoxin reductase 1 (TXNRD1; 2.8 fold increase, 6 hours)^27^. Another cDNA microarray study using androgen-sensitive prostate cancer cells identified a 2.6 fold increase in SOD1 at 10nM of 1,25D_3_ treatment for 24 hours^27^. Calcitriol was also found to induce glucose-6-phosphate dehydrogenase (G6PD) transcription in a VDR-dependent manner within non-malignant prostate cancer cells to maintain NADPH levels for the production the ROS scavenger glutathione^28^. To compliment these studies, the aim of this research was to analyze the potential vitamin D-mediated regulation of the “SOD1-to-SOD2 switch” within the MG-63 p53-independent osteosarcoma model that was recently characterized between non-tumorigenic and tumorigenic breast cancer cells^29^.

## Results

### Effects of 1,25D_3_ on MG-63 osteosarcoma cell viability using the MTT assay

In order to establish the *in vitro* conditions to investigate the mechanism of action toward mitochondrial dynamics, MG-63 cellular response to a 10-fold serial dilution of 1,25D_3_ was assessed using the 3-[4,5-dimethylthiazol-2-yl]-2,5 diphenyl tetrazolium bromide (MTT) assay (**Figure 1**). MG-63 cells were treated with 1,25D_3_ at concentrations ranging from 1pM to 1μM, and cell viability was assessed at 24 and 48 hours post treatment. In addition, two different media conditions were compared to access the synergistic effects of liganded VDR signaling. For one, a charcoal-stripped fetal bovine serum (FBS)-based media was applied to assess the singular role of VDR signaling within MG-63 osteosarcoma cells. For the other, a standard complete media with normal FBS was applied to study the synergistic effects of VDR signaling with other hormone ligands in the system. Interestingly, when cultured with charcoal-stripped media, 1,25D_3_ was unable to inhibit viability of MG-63 cells even at the highest concentration of 1μM for both 24 and 48 hours after treatment (**Figure 1A**). However, in complete media a mild suppression of cell viability was observed only after 24 hours of 1,25D_3_ treatment at the higher concentration range of 10-1000nM (*p*<0.05-0.001) (**Figure 1A**). This suppression in cell viability became even more pronounced after 48 hours of 1,25D_3_ treatment within the lower 0.1-1000nM range (**Figure 1A**; *p*<0.05-0.001). Based on these studies, it appears that 1,25D_3_ treatment over time has a significant suppressive effect on MG-63 cell viability within human serum levels down to the pmol/L range^30^.

**Figure 1.**
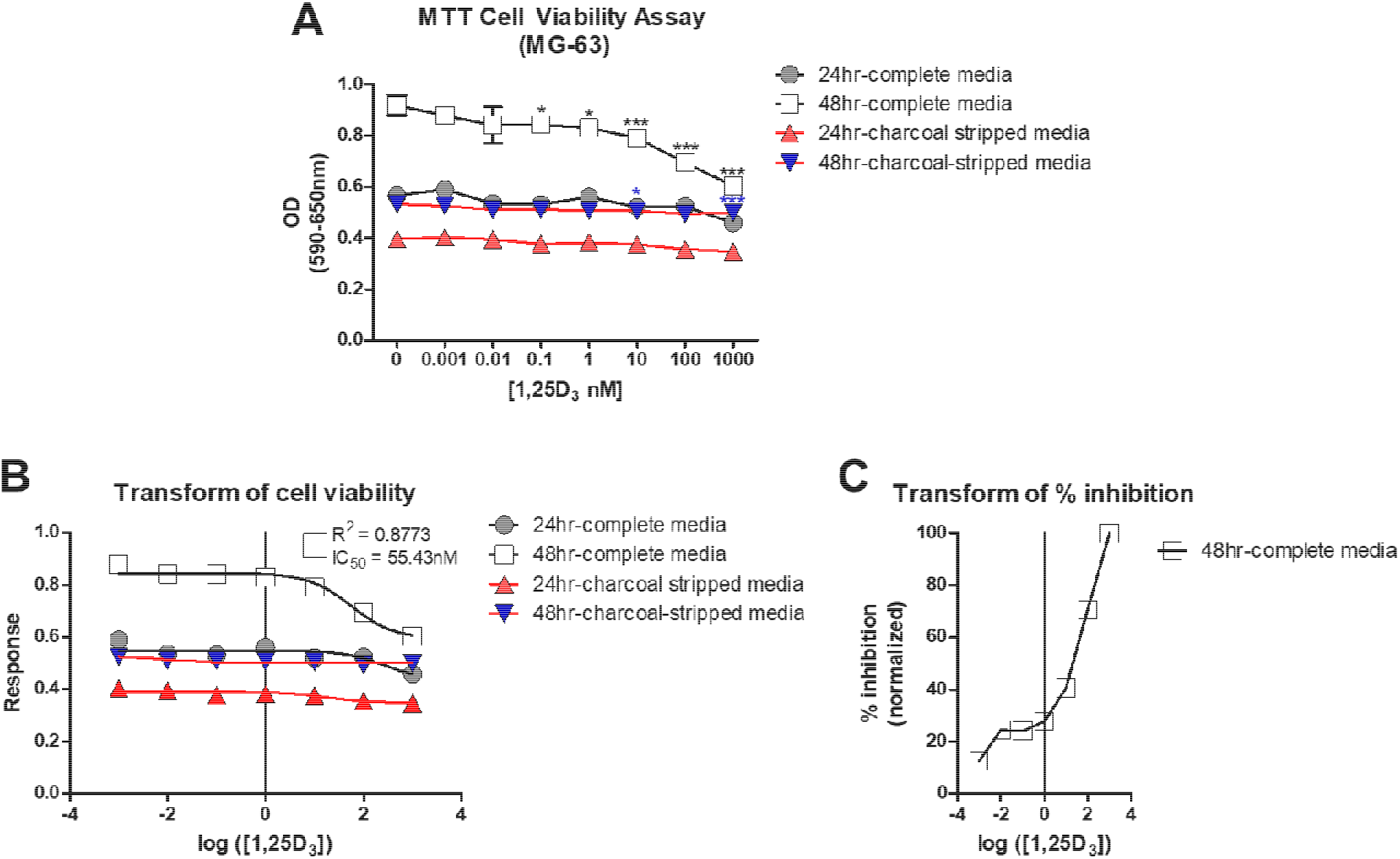
**A) The impact of 1,25D_3_ on MG-63 cell viability using the MTT assay.** The MTT assay was performed in experiments with complete serum or charcoal-stripped serum supplementation for 24 and 48 hours of incubation. The cells were treated with one addition of 1 pM to 1μM 1,25D_3_ before the determination of cell viability. *p*≤ 0.05*, 0.01**, 0.001*** using the 2-way ANOVA with Bonferroni post hoc test (n=5; error bars represent the SD). **B) Logarithmic dose vs. response graph (MTT assay 24-48 hours)**. The calculated logIC_50_ for 1,25D_3_ in 48 hours was 1.74 and the IC_50_ was 55.43nM. In this figure the Hill slope is 1.0, which describes the steepness of the curve. The value R^2^ (R square) quantifies goodness of fit. It is a fraction between 0.0 and 1.0 and has no units. Higher values indicate that the model fits the data better. In this graph, R square is 0.8773 and degree of freedom = 34. **C) Degree of inhibition by 1,25D_3_ on MG-63 osteosarcoma cells after 48 hours of treatment.** The maximal inhibition value was obtained with 1 μM 1,25D_3_ and arbitrarily set to 100%.

Based on our dose curve for MG-63 cells, the IC_50_ value (i.e. the concentration that provides a response halfway between the maximal and the minimally inhibited response) for 1,25D_3_ was determined (**Figure 1A**). The 1,25D_3_ concentrations were transformed to log value, whereby the model assumes that the response curve has a standard slope equal to a Hill slope (or slope factor) of −1.0. As there were no maximal response for the 24-hour complete media and 24/48 hour charcoal-stripped media treated samples, the IC_50_ values could not be determined in these cases. For the 48-hour complete media condition, the best-fit IC_50_ value based on non-linear regression was 55.43nM (logIC_50_ = 1.74), whereby the goodness of fit R^2^ value was 0.8773, and correlated with the transformed % inhibition data (**Figure 1C**). In order to study the involvement and regulatory mechanisms of 1,25D_3_ on mitochondrial dynamics, 10nM and 100nM concentrations were further investigated which encapsulates the determined IC_50_ value.

### Characterization of 1,25D_3_-mediated cell cycle and vitamin D catabolic and osteogenic genes in MG-63 osteosarcoma cells

To acquire further insight into the molecular mechanisms underlying the increased inhibition of MG-63 osteosarcoma cell viability by 1,25D_3_, gene expression analysis was performed of cell cycle, vitamin D signaling and osteogenic-related genes. For this part, MG-63 cells were treated with 10nM and 100nM 1,25D_3_, or vehicle (equi-volume ethanol) for 24 and 48 hours in complete media. First, proliferating cell nuclear antigen (PCNA) mRNA was appraised by real-time PCR to assess the level of DNA replication in the samples. After 1,25D_3_ treatment at 10nM, a statistically significant >2-fold suppression of PCNA transcripts was observed compared to vehicle-treated samples for both the 24- and 48-hour time points (**Figure 2A**). However, no large difference in PCNA transcript levels was observed at the higher 100nM of 1,25D_3_ at both time points, which was also confirmed upon vitamin D antagonist ZK159222 (VAZ) treatment that blocks VDR genomic actions^31^ (**Figure 2A**). These results suggest that 1,25D_3_ suppresses DNA replication (i.e. cell division) only at 10nM in the system.

**Figure 2.**
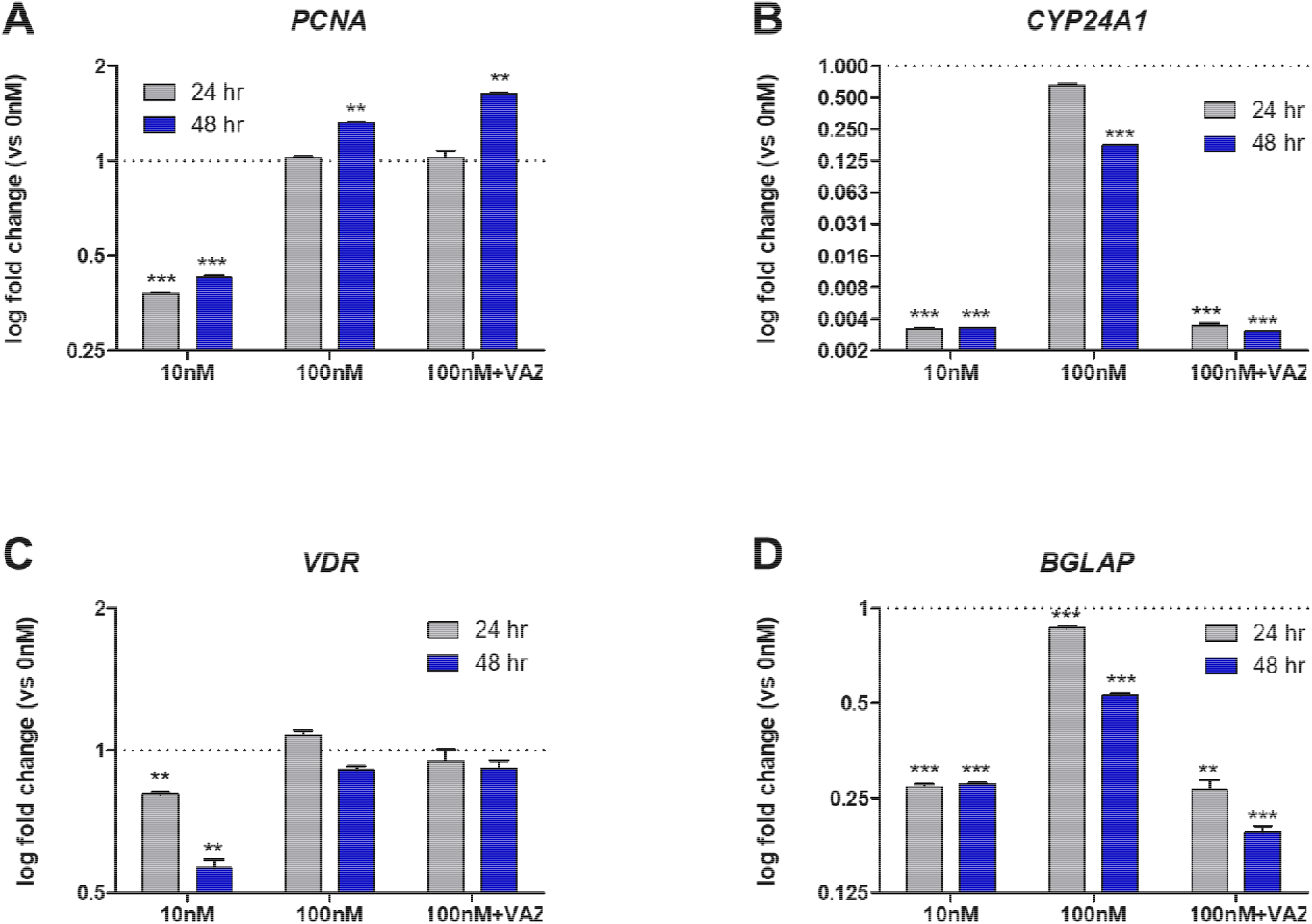
**A) PCNA transcript log-fold response.** Real-time PCR analysis of MG-63 cells in complete media. **B) CYP24A1 transcript log-fold response.** Real-time PCR analysis of MG-63 cells in complete media. **C) VDR transcript log-fold response.** Real-time PCR analysis of MG-63 cells in complete media. **D) BGLAP transcript log-fold response.** Real-time PCR analysis of MG-63 cells in complete media. *p*≤ 0.05*, 0.01**, 0.001*** using the 2-way ANOVA with Bonferroni post hoc test (n=3; error bars represent the SD). VAZ (vitamin D receptor antagonist ZK159222)

In order to monitor the control of vitamin D signaling, cytochrome P450 family 24 subfamily A member 1 (CYP24A1) and VDR transcript levels were monitored. CYP24A1 is a vitamin D_3_ 24-hydroxylase and a VDR target gene that is involved in the catabolism of calcitriol. After treatment of MG-63 cells with 10nM 1,25D_3_ for both 24 and 48 hours, there was a significant reduction in the level of CYP24A1 compared to vehicle-treated controls (**Figure 2B**). However, after 100nM 1,25D_3_ treatment, there was a significant increase in CYP24A1 mRNA compared to 10nM-treated samples. This response was mediated by genomic VDR signaling, as a significant decrease in CYP24A1 mRNA was observed after VAZ co-treatment. This observation suggests a vitamin D-dependency at the 100nM concentration, and an overall suppression of vitamin D signaling through 1,25D_3_ catabolism. VDR mRNA level was measured and shown to be significantly decreased after 10nM 1,25D_3_ treatment that was more striking after 48 hours (**Figure 2C**). However, there was no change in VDR mRNA levels after 100nM 1,25D_3_ treatment at both time points. Furthermore, the VDR mRNA level was not altered after VAZ treatment confirming the presence of receptor in the system and limited autoregulation^7^. Although CYP24A1 is commonly upregulated in cancers and thought to be one mode for vitamin D resistance^32^, this appears not to be the case for MG-63 cells treated with 100nM 1,25D_3_ based on the MTT results (**Figure 1A**). This suggests other anti-cancer strategies upon high levels of vitamin D, including apoptosis^33^ and/or induction of other stress pathways such as those involving the endoplasmic reticulum^25^.

In addition, osteocalcin (encoded by *BGLAP*) mRNA levels were monitored. Osteocalcin is a secreted non-collagenous protein hormone often used to assess osteoblast activity, which is under direct VDR regulation^34^. At 10nM 1,25D_3_ treatment for both 24 and 48 hours, a significant and drastic decrease in BGLAP mRNA levels was observed (**Figure 2D**). However, an increase to near baseline levels of BGLAP mRNA was observed after 100nM 1,25D_3_ treatment, which was abrogated upon VAZ treatment confirming VDR dependency. As increased osteocalcin production is known to cause endoplasmic reticulum stress coupled with elevated oxygen consumption^35^, the differences between 10nM and 100nM 1,25D_3_ may further represent a switch from cytostatic to cytotoxic metabolism, respectively.

### Vitamin D treatment does not inhibit the transcription of respiratory chain genes *COXII* and *COX4I1* in MG-63 cells

Recent studies have shown that vitamin D can suppress the production of cellular ROS in MG-63 cells^25^, however the mode of action is unknown. First mitochondrial and nuclear transcription of cytochrome c oxidase subunit 2 (COXII) and subunit 4 isoform 1 (COX4I1) mRNA, respectively, were characterized after 1,25D_3_ treatment over time as potential players in ROS regulation. These proteins are subunits of cytochrome c oxidase complex IV part of the active mitochondrial respiratory chain and contribute to the proton electrochemical gradient across the inner mitochondrial membrane. Previous studies have shown that both of these subunits were inhibited after 1,25D_3_ treatment in skin keratinocyte models^36^, and may be involved in osteosarcoma. However, after both 10nM and 100nM 1,25D_3_ and VAZ co-treatments of MG-63 cells for 24 and 48 hours, no striking differences in the gene expression of both *COXII* and *COX4I1* were observed (**Figure 3A,B**). These results suggest that 1,25D_3_ does not modulate active cytochrome c oxidase via COXII and COX4I1 subunits at the transcriptional level. These results point to alternative mitochondrial regulators, including perhaps other components of the electron transport chain, in the regulation of ROS by vitamin D^25^.

**Figure 3.**
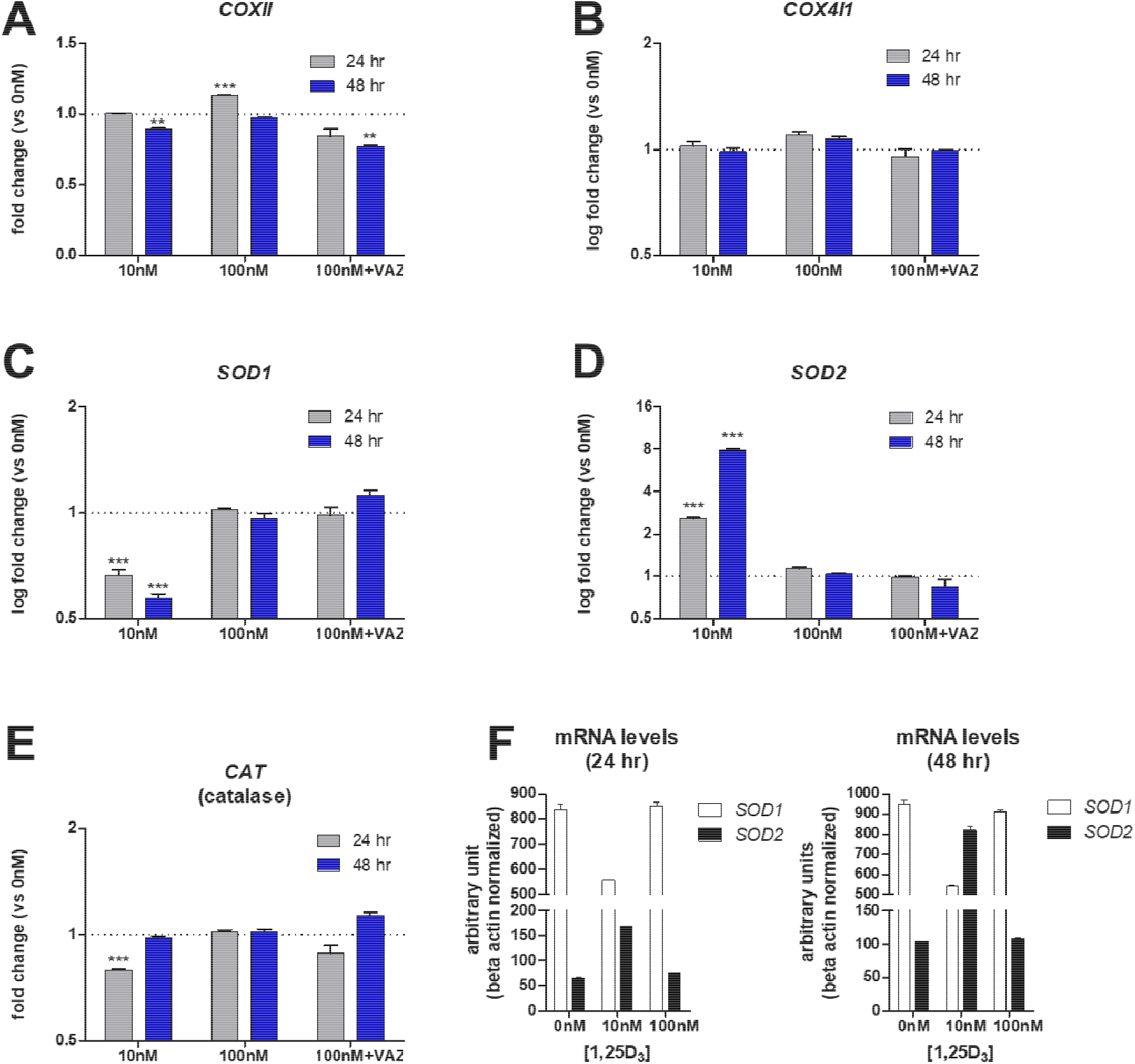
**A) COXII transcript log-fold response.** Real-time PCR analysis of MG-63 cells in complete media. **B) COX4I1 transcript log-fold response.** Real-time PCR analysis of MG-63 cells in complete media. **C) SOD1 transcript log-fold response.** Real-time PCR analysis of MG-63 cells in complete media. **D) SOD2 transcript log-fold response.** Real-time PCR analysis of MG-63 cells in complete media. **E) CAT transcript log-fold response.** Real-time PCR analysis of MG-63 cells in complete media. **F) SOD1-to-SOD2 switch.** Transcript levels presented as arbitrary units normalized to beta actin amounts. *p*≤ 0.05*, 0.01**, 0.001*** using the 2-way ANOVA with Bonferroni post hoc test (n=3; error bars represent the SD). VAZ (vitamin D receptor antagonist ZK159222)

### SOD1-to-SOD2 antioxidative switch after vitamin D treatment of MG-63 osteosarcoma cells

Given the known increase in ROS among MG-63 osteosarcoma cells^25^, it was hypothesized that antioxidant defense mechanisms may be involved in vitamin D-dependent suppression of ROS production. Specific SOD activities are correlated with tumorigenicty^16,29^, therefore the transcription of the SOD genes within 1,25D_3_-treated MG-63 cells was monitored over time. A significant and dramatic decrease in the level of SOD1 transcript was observed only after 24 and 48 hours of 10nM 1,25D_3_ treatment (**Figure 3C**). At 100nM of 1,25D_3_ there was no change in SOD1 levels with or without VAZ, suggesting no direct VDR-dependent regulatory role at higher concentrations of ligand. Interestingly, a dramatic >2.5-8-fold time-dependent persistent increase in SOD2 mRNA levels was observed only with the lower 10nM 1,25D_3_ concentration at both time points (**Figure 3D**). Once again, at the higher 100nM 1,25D_3_ concentration the regulatory role of VDR toward SOD2 activity was abrogated. Catalase is a peroxisomal enzyme that rapidly degrades hydrogen peroxide (H_2_O_2_) to O_2_ and H_2_O. As SODs may yield H_2_O_2_, the mRNA levels of the H_2_O_2_ antioxidant catalase (CAT) were monitored, and a moderate and statistically significant decrease was observed only after 10nM 1,25D_3_ treatment at the 24-hour time point (**Figure 3E**). However, a steady increase following another 24 hours back toward baseline was also observed (**Figure 3E**). This suggests the protective role of vitamin D treatment relying less on CAT antioxidant production during the early stages of treatment, and the potential decrease in H_2_O_2_ efflux from the mitochondria. Lastly, the real-time PCR results were arranged using arbitrary units in order to compare amounts of transcripts across experiments. In this way, it became more evident that after 1,25D_3_ treatment there was a clear shift from SOD1 to SOD2 initiated at the 24-hour time point (**Figure 3F**). However, by the 48hour time point SOD2 levels exceeded SOD1 levels to complete the “switch” (**Figure 3F**). The switch from SOD1 to SOD2 may signify the importance of the cellular distribution and actions of each SOD enzyme, cytosol versus mitochondria, respectively. Overall, these findings suggest an active time and concentration-dependent mitochondrial reparative process involving specific SOD enzymes engaged after vitamin D treatment within MG-63 osteosarcoma cells as part of vitamin D’s anti-cancer effects (**Figure 4**).

**Figure 4.**
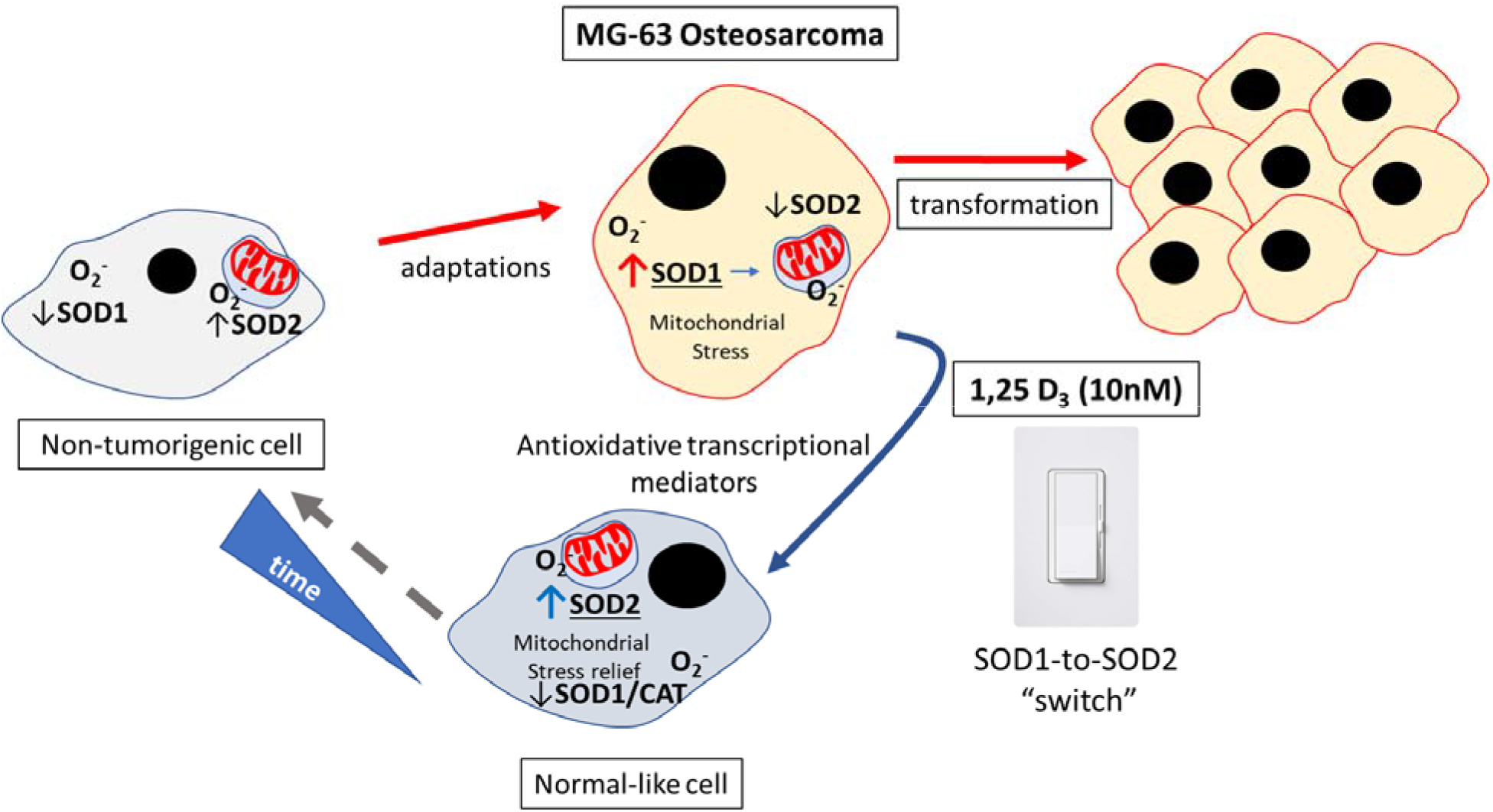
Schema representing the SOD1-to-SOD2 switch in MG-63 osteosarcoma cells after vitamin D treatment.

## Discussion

The major finding of this paper is that MG-63 osteosarcoma cells can respond to adaptive signals such as vitamin D to suppress the transformation process by upregulation of SOD2. The degree to which the vitamin D-SOD network offers specific cytoprotective and antioxidative effects is unclear, and the focus of future studies. Importantly, this form of regulation is tightly controlled in a 1,25D_3_ concentration and time dependent manner which argues for maintenance of proper vitamin D levels in the body. The importance of SODs are reflected in mice either lacking mitochondrial or cytosolic SODs that die around 21 days after birth or exhibit milder phenotypes, respectively, such as the development of cancer in time^37^. It appears if SOD levels and distributions are unchecked, this can contribute to mitochondrial damage and eventual cancer transformation^16^. Recent studies using both human and murine cancer models indicate that the expression of SOD2 is rapidly reduced upon activation of various oncogenes^29^, thus compromising it’s cytoprotective and antioxidative roles, presumably within the mitochondria. Thus, strategies to induce SOD2 may be an effective way to limit cancer transformation. On the other hand, SOD1 is overexpressed in a majority of mammary tumor models driven by various oncogenes^29^. It is possible that mitochondrial SOD1 may be necessary to maintain the integrity of the organelle when the levels of SOD2 are compromised during transformation. Indeed, after 10nM 1,25D_3_ treatment for 48 hours there was a cumulative increase in the combined SOD1 and SOD2 transcript amounts compared to untreated MG-63 cells (**Figure 3F**). Nevertheless, the consensus is that in non-tumorigenic cells SOD2 levels are elevated while SOD1 levels are kept low^29^, suggesting that mitochondrial protection from ROS is critical in the natural high respiratory state. It was recently shown that pharmacological inhibition of SOD1 results in fragmented and dilated cristae of mitochondria in a breast cancer cell line (MDA-MB-231)^29^, suggesting the critical role of SOD1 protection from mitochondrial ROS toxic effects during transformation. However, SOD1 inhibition within non-tumorigenic cells did not alter morphological features of the mitochondria^29^, presumably due to protection from the known increased endogenous levels of SOD2. These findings correlate with the long recognized understanding that cancer cells are characterized by elevated levels of ROS ^38^. Furthermore, it has been reported that SIRT3 – a major regulator of SOD2 - expression is decreased in 87% of breast cancers implying a key role of SOD2 in cancer transformation ^39^. Based on the findings presented here, vitamin D-mediate protection from ROS-induced damage can be attributed to the expression of the ROS detoxifier SOD2. Importantly, it appears that vitamin D has the potential to shift the SOD profile from one that represents a tumorigenic towards a non-tumorigenic state.

It is unclear how *SOD1* and *SOD2* expression is regulated by vitamin D and its receptor through *cis* and/or *trans* regulatory elements and factors. It is clear from the MTT data that other serum ligands work in synergy to support vitamin D’s anti-cancer effects (**Figure 1A**), including the SOD1-to-SOD2 shift. It is possible that vitamin D regulates SOD1 and SOD2 expression via indirect mechanisms. For example, vitamin D is known to activate NRF2, which is a transcription factor that plays an important role in redox homeostasis and protection against oxidative damage^40^. Upon induction, NRF2 translocate to the nucleus and binds DNA antioxidant response elements (AREs) that leads to coordinated activation of gene expression^41^. Thus, vitamin D signaling may promote expression of NRF2 that then regulates antioxidant genes such as *SOD1* and *SOD2,* which remains to be tested^41^.

Previous studies using either *VDR* ablation or 1,25D_3_ treatment approaches have made associations between VDR actions and mitochondrial respiratory activity along with production of intracellular ROS within several cell lines^42^. In all cell types it was found that upon *VDR* ablation there was an increase in respiration and ROS production associated with elevations in both mitochondrial COX2 and nuclear COX4I1^42^. On the other hand, 1,25D_3_ treatment of skin keratinocytes resulted in decreased COX2 and COX4I1 mRNA levels, suggesting a concomitant reduction in intracellular ROS production^36^. Although in this study, evidence was provided for a mitochondrial-specific antioxidant effect of vitamin D within MG-63 cells, no change in either COX2 or COX4I1 transcript levels was found (**Figure 3A,B**). This would suggest that within MG-63 cells other vitamin D-dependent electron transport chain enzyme complexes may be involved, as ROS are sourced from Complex I and III enzymes, while Complex IV converts molecular oxygen to two molecules of water and activates ATP synthase to synthesize ATP^43^. It is well-known that the maintenance of proper vitamin D levels for the long term will have beneficial cancer survival effects across demographics^15^, and in this paper, vitamin D was shown to significantly induce mitochondrial SOD2, and presume to be part of the antioxidant mechanism to limit the proliferation and cellular damage of MG-63 osteoblastoma cells. Future studies will focus on other non-calcemic vitamin D analogues and their effects on the SOD1-to-SOD2 switch and their role in cancer biology.

## Experimental procedures

### Reagents and cell culture

Crystalline 1,25(OH)_2_D_3_ (Biomol, Plymouth Meeting, PA, USA) was reconstituted in ethanol and kept at −80°C. The vitamin D receptor antagonist ZK159222 (VAZ, Toronto Research Chemicals) was reconstituted in ethanol and kept at −80°C (Ochiai E et al. 2005).

Human MG-63 osteosarcoma cells (CRL-1427; American Type Culture Collection, Manassas, VA, USA) were cultured in complete media containing Eagle’s Minimum Essential Medium (ATCC, 30-2003), 10% heat inactivated fetal bovine serum, and 1X penicillin and streptomycin. For some experiments, the cells were cultured in charcoal-stripped FBS (35-072-CV, Corning). For assays, cells were treated with 0 (vehicle) to 1μM 1,25(OH)_2_D_3_ incubated in tissue culture plates (CytoOne).

### Quantitative real-time RT-PCR (qPCR) analyses

RNA was prepared using the PureLink RNA Mini kit (ThermoFisher Scientific). cDNA was synthesized using 200 ng total RNA with the ProtoScript^®^ First Strand cDNA Synthesis kit (New England Biolabs) utilizing random hexamers. All cDNAs were amplified under the following conditions: 95°C for 10 minutes to activate AmpliTaq Gold^®^ Polymerase; followed by 40 cycles of 95°C for 15 seconds and 60°C for 1 minute with an internal ROX reference dye.

qPCR analysis was performed on a QuantStudio 3 Real-Time instrument (ThermoFisher Scientific) utilizing the Power SYBR™ Green PCR Master mix (ThermoFisher Scientific; primer list, Supplemental Table S1). Target genes were normalized to beta actin mRNA expression. For the primer design, the human genome sequence coverage assembly GRCh38.p13 was utilized from the Genome Reference Consortium.

### MTT assay protocol for measuring cell viability

All experiments were performed using 2.4×10^4^ cells/well in 96-well plates. Cells were seeded and then left to incubate at 37°C (5%CO2) overnight. The following day, cells were carefully washed with 1X PBS, and then treated with 1,25D_3_ and the cellular status was assessed at both 24 and 48 hours post treatment. The MTT (3-[4,5-dimethylthiazol-2-yl]-2,5 diphenyl tetrazolium bromide) assay was performed according to the manufacturer’s recommendation (10009365, Cayman Chemicals). After incubation with the MTT reagent, the cells and salt were solubilized using the provided sodium dodecyl sulfate-based lysis buffer. The optical density absorbance was determined using a Molecular Devices EMax microplate spectrophotometer at 550nm absorbance minus a 650nm reference. Data is represented as the mean of 5 replicate wells ± SD, with analysis using the two-way ANOVA with Bonferroni post-hoc test (GraphPad Prism). In order to approximate the IC_50_ value, GraphPad was used and applied nonlinear regression after log transformation of the 1,25D_3_ dose assessed by the model: Y=Bottom + (Top-Bottom)/(1+10^((X-LogIC_50_))).

### Real-time PCR Data analysis

Data are presented as fold induction of treatments compared to 0nM (vehicle) samples normalized to beta actin mRNA levels (i.e. the comparative CT Livak method (Livak, Schmittgen 2001)^44^. Melting curve analysis was performed for all primers to eliminate those that yielded primer-dimers. The *p*-values reflect the log fold-change compared to the vehicle (0nM) condition (*n*=3 experimental samples ± SD). As the delta Ct (dCT) values are measures that are proportional to log expression, a t-test using two groups of dCT values was used to generate the *p*-values.

## Author contributions

T.S.L. designed and performed the experiments, analyzed the data, and wrote the manuscript.

## Acknowledgement

Supported by Grant # IRG-17-183-16 from the American Cancer Society, and from the Sylvester Comprehensive Cancer Center at the Miller School of Medicine, University of Miami

## Data availability statement

All data generated during and/or analyzed during the current study are available from the corresponding author on reasonable request.

## Tables

**Supplemental Table S1.**
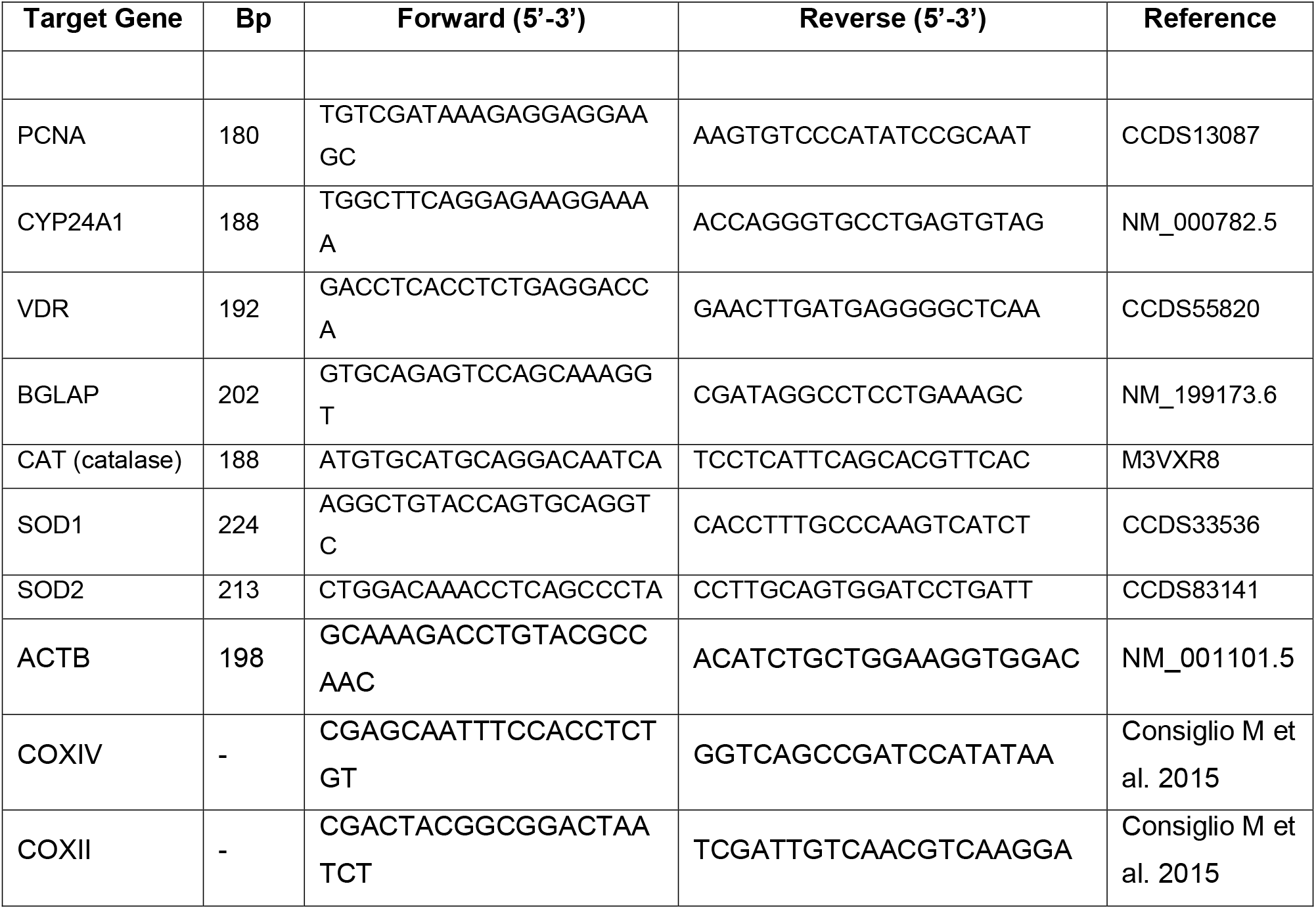
Real-time PCR primer set (human)

